# Rethinking the Risk of Uncertainty: Human-AI Interaction in Household Mycology

**DOI:** 10.64898/2026.02.24.707810

**Authors:** Nik V. Kuznetsov

**Affiliations:** ASPIRE Precision Medicine Research Institute Abu Dhabi, United Arab Emirates University, P.O. Box 15551, United Arab Emirates; Karolinska Institutet, Nobels väg 16, SE-17177 Stockholm, Sweden

**Keywords:** artificial intelligence, AI application, human-AI interaction, computer vision, safety, toxicity, sustainability, species identification, fungi, Macromycetes, mushroom foraging

## Abstract

**Background:** In recent years, the rise of AI-mediated technological assistance has impacted applied mycology. Various tools employ AI to analyze images of Macromycetes fruiting bodies for species identification. This trend has sparked widespread interest in online applications despite the potential risks associated with relying on AI-generated advice.

**Methods:** We conducted a comparative analysis of popular AI-based mushroom identification tools using over 100 original photographs of fungi fruiting bodies from nearly 60 species, taken in real-world conditions. Reference searches were conducted with mushroom names in five languages, including Latin species names. Functional test scores and an overall accuracy score were calculated for twelve selected AI applications to evaluate their general reliability.

**Results:** The AI-based applications evaluated in the study were able to recognize only a portion of the provided mushroom images. Even the best-performing tools frequently failed to accurately identify fruiting bodies in real-world conditions. None of the tested applications consistently provided a single, correct species name. Instead, users were often presented with multiple options, among which the correct answer might have been found.

**Conclusion:** When addressing mycological queries, it is crucial to recognize the inherent risk of relying solely on AI-mediated resources for mushroom identification. Various limitations hinder their effectiveness in real-world environments. These tools should only be viewed as supplementary aids since they are inadequate for making definitive or safety-critical decisions.

## Introduction

Science fiction often portrays apocalyptic scenarios in which computers rebel against humanity, culminating in the rise of a militant AI and eventual destruction of civilization. While these scenarios remain firmly in the realm of fiction, it is worth noting that even imperfect AI systems, prone to errors, can pose risks comparable to those of rogue or malicious AI.

The rapid proliferation of AI technologies across nearly every domain of human cultural and daily life, while yielding significant advancements, also introduces notable risks [1-3]. These risks are particularly pronounced when the users place unwarranted trusts in AI as a reliable source of information or disregard warnings about its possible inaccuracies. A clear example of this trend is the proliferation of apps that claim to provide “guaranteed” identification of natural objects.

Broadly, users of these AI applications in natural settings can be categorized into three groups, each facing varying levels of risk when making crucial decisions influenced by AI-generated guidance. Recreational activities, such as exploring wildlife and identifying plants or fungi during forest walks, generally pose minimal risks. Mushroom foraging, has deep roots in cultural traditions and has generates scientific interest in fungal communities and their ecological interactions [4-8]. When these applications are used for scientific purposes requiring precise species identification, AI errors can lead to inaccuracies that may compromise research outcomes.

The risks become more consequential when mushrooms are identified for collection and consumption. Mistaken identification and decisions regarding edibility can result in illness or, in severe cases, have fatal outcomes. Regrettably, AI-mediated errors in mushroom identification have already been documented, with an increasing number of incidents reported [9,10]. As the old Irish adage wisely warns: “There are old mushroom hunters and there are bold mushroom hunters. But there are no old bold mushroom hunters.”

Below, we outline the top ten primary complications and causative factors contributing to the malfunctions of AI applications. Additionally, 12 AI-based tools (six software programs and six mobile applications) were evaluated using a scoring system based on these factors. An overall score was then calculated to assess the general accuracy and practical utility of each application.

## Methods

Over the course of ten years, digital photographs of fruiting bodies of macromycetes were collected from various forested regions and local markets across Eurasia. European species were identified using established field identification handbooks in combination with the researchers’ extensive foraging experience. Asian species were identified in collaboration with a mycology expert possessing over 25 years of taxonomic expertise. In total, 103 high-quality digital photographs were selected, representing 62 species across 40 genera. For analysis, 48 representative images were organized into seven thematic Topic Test Groups (TTGs). Identification queries were submitted using Latin species names as well as common names in four languages: English, Swedish, Russian, and Chinese (Supplementary Table 1).

An initial screening of over 30 applications and websites was conducted based on publicly available user ratings from the Apple App Store, Google Play Store, and Google search engine results. A total of twelve publicly available Fungi Identification AI-based Resources (FIARs) were selected for comparative analysis, comprising six mobile applications and six web-based platforms (see Figure 1; Supplementary Figure 1; Supplementary Table 2). These tools were evaluated side-by-side to assess their performance under real-world conditions. The total score, or Total Rating Score (TRS), was calculated for each application across all tests (Supplementary Table 2).

**Figure 1.**
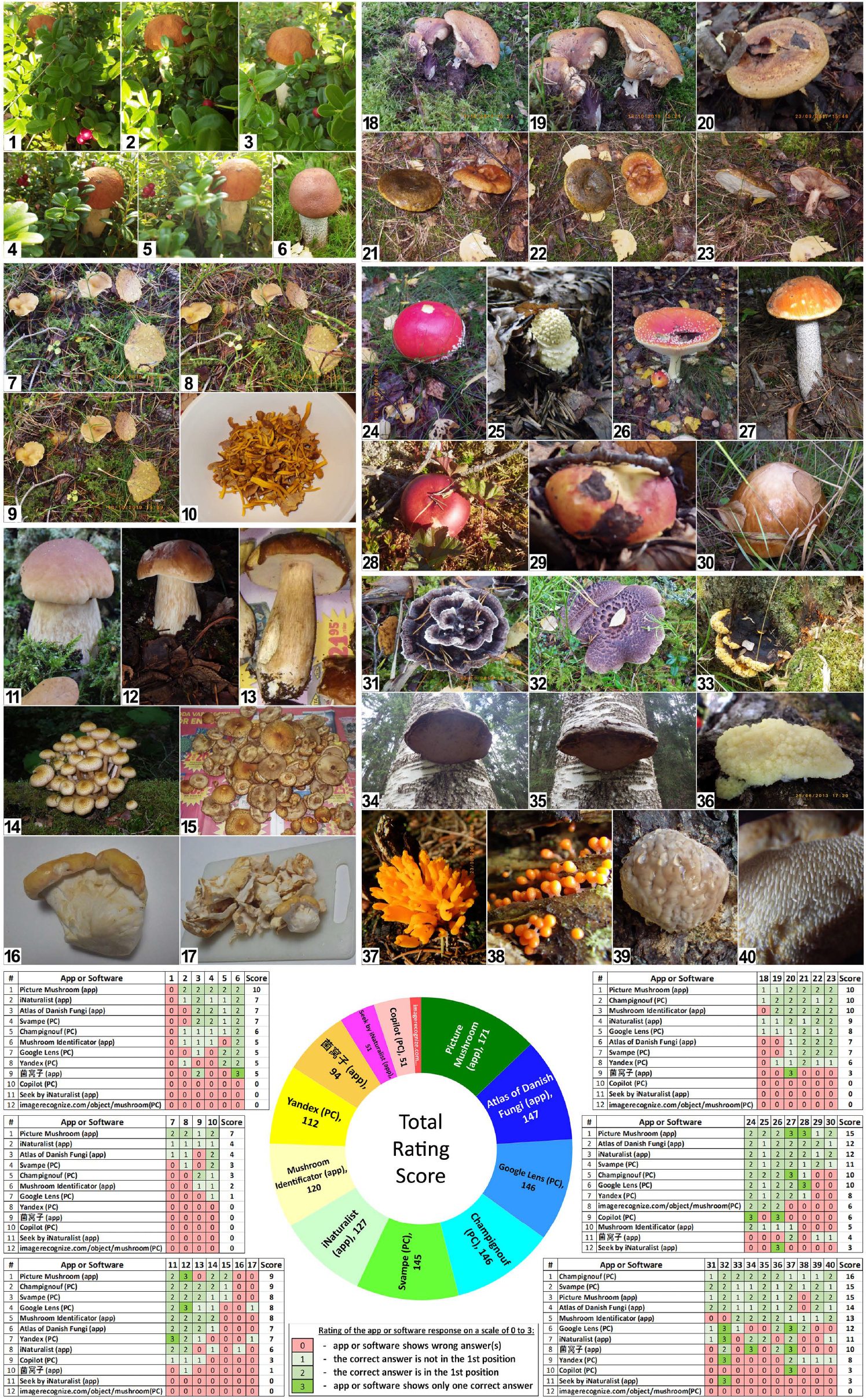
Comparative analysis of Fungi Identification AI-based Resources. A comparative analysis of twelve Fungi Identification AI-based Resources (FIARs) was performed, using a set of real-world photographs. The doughnut chart illustrates Total Rating Score (TRS), calculated to assess the general accuracy and practical utility of each AI tool (see text for details).

Three researchers independently tested the 12 selected FIARs using personal mobile devices and computers. Each researcher reviewed the full list of AI-generated responses for each query, recorded the correct identification (if present), and noted its position in the list. The devices used were as follows: Researcher 1 operated an iPhone™ 13 (Apple Inc., 2021); Researcher 2 used a Motorola G54 smartphone (Motorola Mobility, 2023); Researcher 3 used a Samsung Galaxy Tab S8 (Samsung Electronics, 2022) and PC notebook Vivobook 16X (M1603) (Windows 11) (ASUS, 2022).

The responses of each AI application were evaluated using a four-point scale (0-3) as follows: 0 – only incorrect answers listed; 1 - the correct answer included but not ranked first; 2 - the correct answer ranked first among suggestions; 3 - only the correct answer provided.

To assess overall performance, a Total Rating Score (TRS) was calculated for each FIAR by summing its scores using Microsoft Excel (Baton Rouge, LA, USA). These cumulative results are illustrated in the doughnut chart (Figure 1), providing a comparative overview of general reliability and practical utility across all 12 tested platforms. Additional localized scores were compiled by scenario for each FIAR within each TTG and visualized in the Excel tables embedded in Figure 1 and Supplementary Figure 1, reflecting the outcomes of the ten major complication categories:

1. “Occlusion”
2. “Camouflage”
3. “Influence of lighting, contrast, and background”
4. “Processed mushrooms”
5. “Mushroom morphology variation”
6. “Multiple species per image”
7. “Interspecific coloration similarity”
8. “Basidiomycete species diversity”
9. “Mislabeling and misinformation”
10. “Geographical and temporal biases”

## Results and Discussion

### 1. Occlusion

Occlusion is a fundamental challenge in computer vision that arises when an object is partially obscured by another object or its context. For example, a vivid red-capped scaber stalk (*Leccinum aurantiacum)* occluded by lingonberry shrubs in the first image (Figure 1 – Photo #1) was not correctly identified by any of 12 applications tested (Figure 1; Supplementary Table 1). Yandex, for instance, recognized only the lingonberry leaves, while several other AI tools misidentified the object. Copilot suggested *Boletus reticulatus*, Champignouf identified *Psathyrella candolleana*, and Danish software Svampe purposed the term “Dear wax hat,” whose meaning remains unclear. The Google Lens application missed the mushroom entirely, focusing instead on the red spots of lingonberry leaves in the foreground. Consequently, it identified the species as *Exobasidium rhododendri*, a basidiomycete closely related to the redleaf fungus (*Exobasidium vaccinii)*, which causes galls on lingonberry leaves [11].

The partially occluded mushrooms in Photos #2, 3, 4, and 5 were recognized by a total of five, eight, six, and seven different applications, respectively. However, only a few of these applications correctly identified the species. In contrast, 9 out of the 12 applications successfully recognized and correctly identified the species of the non-occluded mushroom in Photo #6.

Even minor occlusion caused by contextual elements contributed to the misidentification of several other fungal genera and species by many of the AI applications (Photos #28 -30).

### 2. Camouflage of Mushrooms

Many northern fungal species produce fruit bodies during autumn that exhibit dull, earth-toned coloration, blending into the surrounding forest [12,13]. The cryptic appearance, or crypsis, likely provides protection by reducing the likelihood of detection by animals prior to spore dispersal [14,15]. While mimicry in fungi remains a debated topic due to limited empirical evidence [12,16,17], some species exhibit highly effective camouflage that can confuse both humans and AI systems.

A notable example is the yellowfoot or winter chanterelle *(Craterellus tubaeformis)*, whose fruiting bodies are difficult to detect in the deciduous forests, even for experienced mushroom foragers. Unsurprisingly, many AI applications fail to identify this species in natural settings (Photos #7-9). Remarkably, some AI tools, including Copilot, did not recognize it even after it had been cleaned and placed in a domestic setting, such as on a kitchen counter (Photo #10).

### 3. Influence of Lighting, Contrast, and Background

In image-based object recognition, contrast and background are critical variables—and fungal identification by AI tools is no exception. In general, higher image contrast is associated with improved recognition accuracy. For example, a photo of *Boletus edulis* taken at night with flash against a dark background (Photo #12) produced better AI recognition results than a similar image taken during the day under diffuse light with a green moss background (Photo #11), or one photographed under artificial lighting on a colored newspaper (Photo #13). These findings underscore the importance of lighting conditions, image contrast, and background uniformity in AI-mediated species identification.

Similar trends were observed with other fungal species. *Pholiota squarrosa*, was recognized by 7 out of 12 applications when shown in a high-contrast image with a natural background (Photo #14), compared to only 3 out of 12 applications when photographed under artificial lighting with a colored, non-natural background (Photo #15). This further illustrates the importance of background context and lighting quality in AI-mediated identification.

### 4. Processed Mushrooms

As anticipated, photographs of collected and processed mushrooms proved more difficult for AI systems to identify accurately. A previous study using real-world images submitted by the public in Australia reported similar findings [18]. Photos of mushrooms prepared for cooking often featured incomplete specimens, suboptimal lighting, and non-standardized backgrounds, all of which negatively impacted recognition performance.

Similar misidentification issues were observed for several commonly foraged European species when presented in processed form. Notable examples include images of *Boletus edulis* (Photo #13), *Pholiota squarrosa* (Photo #15 vs. Photo #14), *Albatrellus ovinus* (Photos #16, #17) at different preparation conditions. These cases highlight how the physical alteration of specimens can obscure key morphological features necessary for accurate AI-based identification.

### 5. Mushroom Morphology Variation

Basidiomycetes are known for their considerable intraspecific morphological variation [19,20]. Most AI applications, predictably, are trained and biased toward recognizing fungi with “standard” or typical appearances. The variation of fungi fruiting bodies poses a considerable challenge for accurate identification. For instance, a group of three fruiting bodies of *Paxillus involutus* exhibiting atypical morphology (Photos #18, #19), markedly different from the classic form (Photo #20), were not recognized by 6 to 8 of the tested applications.

Moreover, morphological differences between the earlier stages and its mature fruiting bodies can substantially reduce the accuracy and reliability of AI-based identification. A well know example is the toxic fly agaric (*Amanita muscaria*), whose “button” stage appears distinctively different from its mature form [21]. This disparity often leads to misidentification, particularly among inexperienced mushroom foragers. While it may be expected that AI systems could resolve such challenges, current tools consistently fail to identify early-stage specimens with acceptable accuracy.

Particular caution is warranted when using AI tools to identify immature mushrooms. Imaging of fruiting bodies at early developmental stages presents substantial challenges for species recognition. In this study, only 3 of the 12 tested AI applications successfully identified *Amanita muscaria* at its “button” stage (Photo #25), compared to 10 of 12 applications that correctly identified the species in its mature form (Photos #24, #26).

### 6. Imaging of Multiple Species in a Single Photo

The presence of multiple fungal species within the same image complicates the identification process, as many AI tools appear to be designed to recognize only a single species per photograph. For instance, when *Paxillus involutus* appeared alone (Photo #20), the Chinese application accurately identified it. However, the same application failed to recognize *P. involutus* when it was pictured alongside *Lactarius turpis* (Photos #21, #22, #23), highlighting the limitations of multi-species recognition in current AI systems.

Camera angle also influenced the ability of AI tools to accurately identify multiple species within a single image. Notably, Yandex performed better when the fungi were photographed from an elevated angle (Photo #21), while iNaturalist produced more accurate results from a lower and direct top-down presentation (Photos #22, #23). Picture Mushroom, Mushroom Identificator, Atlas of Danish Fungi, and Svampe demonstrated consistent identification performance across different camera angles.

### 7. Similarity of Interspecific Coloration

Visual similarity in coloration and surface patterns across different fungi species present significant challenges for AI-based identification. For instance, the orange-colored cap of *Leccinum aurantiacum* with white specks (Photo #27) was incorrectly classified by Copilot as *Omphalotus olearius* (Jack-O’Lantern), *Mycena leaiana* (orange mycena), or *Amanita muscaria var. guessowii* (orange fly agaric).

In another case, Russula sp. (Photo #29) was misidentified by Champignouf application as *Baorangia bicolor, Leratiomyces ceres, Imperator sp*., *Boletus edulis*, or *Amanita muscaria*. One AI tool (imagerecognize.com) even misclassified the specimen as an apple, with confidence of 99 %.

Compounding the issue, environmental factors can alter key features. For example, the white scales typical of mature *Amanita muscaria* can be washed off in wet weather conditions (Photo #24), increasing the risk of confusion with red *Russula* species. Similarly, *Leccinum scabrum* (birch bolete) was misidentified by Copilot and Google Lens as *Boletus edulis*, and further confused with *Suillus granulatus* (weeping bolete) by Yandex and Champignouf, particularly when the photograph was taken in damp conditions (Photo #30).

### 8. Species Diversity in Basidiomycetes

In addition to morphological variability, the vast taxonomic diversity within the Division Basidiomycota presents a substantial challenge for accurate species identification. Even among experts, classification remains unsettled for many taxa. A significant portion of the time at mycological conferences, including the recent International Mycological Congress (IMC12, 2024), is often devoted to resolving the systematic positions and updating nomenclature.

A striking example is the former classification of the genus *Armillaria*, which historically served as a “wastebasket” taxon encompassing over 270 species of agaric mushrooms with similar macroscopic traits, such as stem-attached gills, a ring, and a white spore print. Following taxonomic revision in 1995, only about 40 species remained within *Armillaria*, while the others were reassigned to 43 separate genera [22].

Many fungal species cannot be reliably identified by appearance alone, requiring DNA sequencing and phylogenetic analysis for accurate classification. This poses a limitation for AI tools, which typically rely on visual data. Furthermore, species with limited culinary, ecological, or medicinal relevance are often underrepresented in training datasets, resulting in poor recognition by AI applications biased toward commonly known edible or toxic fungi (Photos #31 - #40).

### 9. Mislabeling and Misinformation

Accurate labeling is critical for the reliability of AI systems, particularly in tasks involving species identification. Mislabeling by non-specialists, such as photo authors or content uploaders, can introduce noise into datasets and misguide the training and performance of AI models.

A common example involves images of wood-decaying fungi such as *Fomes fomentarius, Phellinus igniarius*, and *Fomitopsis pinicola*, which are frequently mislabeled online as “chaga” (чага). However, true chaga refers specifically to the sclerotium of *Inonotus obliquus*, a distinct species used in folk medicine and subject to ongoing taxonomic and pharmacological debate [23–27].

Additionally, AI systems themselves contribute to the issue by arbitrarily assigning incorrect species names to images, sometimes even generating fictitious mushrooms and labeling them with legitimate scientific names [28,29]. Such errors degrade the overall accuracy and trustworthiness of AI-generated identifications.

### 10. Geographical and Temporal Biases

Although many modern AI platforms are globally accessible, their performance often reflects the geographical origin and temporal scope of their training datasets. As a result, certain tools exhibit regional biases that limit their effectiveness outside of their primary cultural or ecological context.

For example, the yellowfoot or winter chanterelle (*Craterellus tubaeformis*), known as *Trattkantarell* in Scandinavia, is a common and widely foraged mushroom in Northern Europe. However, it was not recognized by regionally developed platforms such as Yandex (Russia) and 菌窝子 (China). Similarly, species widely consumed in Western countries, *Boletus edulis* (porcino), *Leccinum scabrum* (birch bolete), *Suillus granulatus* (weeping bolete) were not identified by 菌窝子.

Conversely, mushrooms familiar in Eastern markets, such as termite mushroom, 鸡枞菌(*Termitomyces yunnanensis*) [30], flat chanterelle, 鸡油菌 (*Cantharellus applanatus*) [31], and rugiboletus, 皱牛肝菌 (*Rugiboletus extremiorientalis*) [32], were not reliably recognized by AI applications primarily developed for Western use.

Among the tested platforms, the Chinese application 菌窝子 demonstrated superior performance with Eastern species (Supplementary Figure 1 – Supplementary Photos #1 - 8), but even this app failed to identify some novel fungi. For example, matijun (馬蹄均, *Turbinellus matijun, Gomphus matijun*), a recently introduced edible mushroom in Asian markets [33], was not recognized by any of the 12 tested AI platforms. This highlights the limitations of static training datasets and the need for regular updates to accommodate emerging species.

## Conclusion

Identifying mushrooms based solely on visual appearance is inherently challenging, even for trained experts. While AI-powered platforms continue to grow in popularity, their effectiveness in real-world scenarios remains limited. All of the applications evaluated in this study showed only conditional accuracy and reliability and failed to identify several fungi images, each demonstrating some degree of limitation. As such, users should exercise caution when relying on these tools, particularly when identification carries potential health or food safety risks. Correct fungal identification via AI is not guaranteed and, ultimately, remains the responsibility of the user.

The use of AI in natural object identification, such as fungal species, represents only a small example of the broader concerns tied to artificial intelligence. Many risks—both current and future—are poorly defined and often underestimated. As human–AI interaction becomes increasingly ubiquitous, it is important to question not only the capabilities of AI, but also the assumptions we make about our role in guiding its use.

## Abbreviations

The following abbreviations are used in this manuscript:

AI: Artificial Intelligence
FIAR: Fungi Identification AI-based Resource
HAI: Human-AI Interaction
TRS: Total Rating Score
TTG: Topic Test Groups

## Author Contributions

N.V.K. conceptualized and designed the study, acquired the data and conducted analyses, wrote the manuscript and prepared the figures. Author approved the submitted version.

## Acknowledgments

The author thank the IMC12 Congress Organizing Committee for interesting discussions and Prof. Dr. Zhu-Liang Yang at the Kunming Institute of Botany (Chinese Academy of Sciences) for invaluable assistance.

## Funding

The study was supported by the ASPIRE Precision Medicine Institute in Abu Dhabi (fund code 21R098).

## Conflicts of Interest

The author declares no conflict of interest.

## Data availability statement

All data supporting the findings of this study are available within the paper and its Supplementary Information.

## Supplementary files

**Supplementary Figure 1. Asian markets fungi species were poorly identified by Western FIARs** Mushrooms, beloved in East, present a challenge to the versatility of AI-based applications intended primarily for Western use. Mushrooms from Asian markets are more effectively recognized by the Chinese app.

**Supplementary Table 1: KEY Reference Table - Mushroom picture numbers and Names**

## Notes

### Competing Interest Statement

The authors have declared no competing interest.

