## Supplementary Figure 1 for "Rethinking the Risk of Uncertainty: Human-AI Interaction in Household Mycology"

**Asian markets fungi species were poorly identified by Western FIARs**
Mushrooms, beloved in East, present a challenge to the versatility of AI-based applications intended primarily for Western use. Mushrooms from Asian markets are more effectively recognized by the Chinese app. The table within the figure was created in Microsoft Excel.

Rating of the application response on a scale of **0** to **3**:

| | | |
|---|---|---|
| App or software shows wrong answer(s) | - | 0 |
| The correct answer is not in the 1st position | - | 1 |
| The correct answer is in the 1st position | - | 2 |
| Application shows only one correct answer | - | 3 |

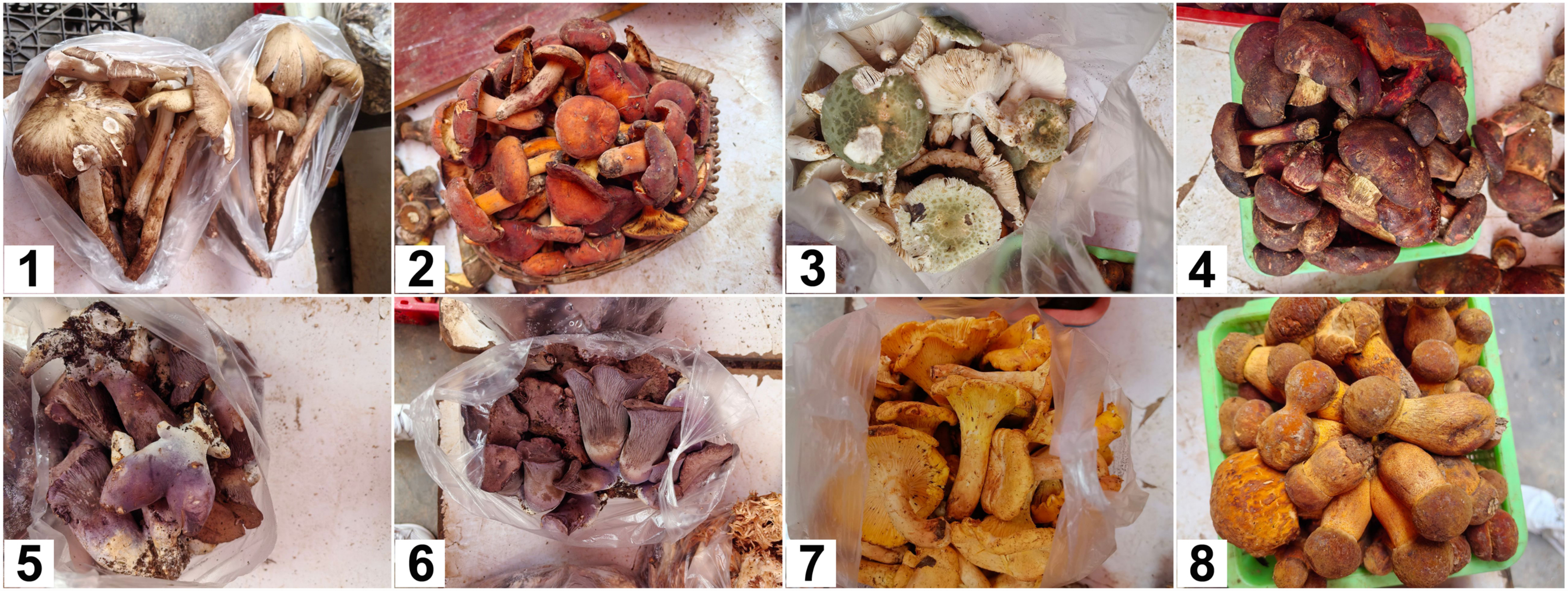


| # | App or Software | Image Number | | | | | | | | Score |
|---|---|---|---|---|---|---|---|---|---|---|
| | | 1 | 2 | 3 | 4 | 5 | 6 | 7 | 8 | |
| 1 | 菌窝子 (app) | 3 | 3 | 3 | 0 | 0 | 0 | 2 | 0 | **11** |
| 2 | Atlas of Danish Fungi (app) | 0 | 1 | 2 | 2 | 0 | 0 | 0 | 0 | **5** |
| 3 | Google Lens (PC) | 1 | 2 | 2 | 0 | 0 | 0 | 0 | 0 | **5** |
| 4 | Picture Mushroom (app) | 0 | 2 | 2 | 0 | 0 | 0 | 0 | 1 | **5** |
| 5 | iNaturalist (app) | 0 | 2 | 1 | 0 | 0 | 0 | 0 | 1 | **4** |
| 6 | Champignouf (PC) | 0 | 0 | 2 | 1 | 0 | 0 | 0 | 0 | **3** |
| 7 | Svampe (PC) | 0 | 1 | 2 | 0 | 0 | 0 | 0 | 0 | **3** |
| 8 | Yandex (PC) | 0 | 0 | 0 | 0 | 0 | 0 | 0 | 2 | **2** |
| 9 | Mushroom Identificator (app) | 0 | 0 | 0 | 0 | 0 | 0 | 0 | 0 | **0** |
| 10 | Copilot (PC) | 0 | 0 | 0 | 0 | 0 | 0 | 0 | 0 | **0** |
| 11 | Seek by iNaturalist (app) | 0 | 0 | 0 | 0 | 0 | 0 | 0 | 0 | **0** |
| 12 | imagerecognize.com/object/mushroom(PC) | 0 | 0 | 0 | 0 | 0 | 0 | 0 | 0 | **0** |
