## Supplementary Table 1 for "Rethinking the Risk of Uncertainty: Human-AI Interaction in Household Mycology"

### Mushroom picture numbers and names

| Image # | Latin | English | Swedish | Russian | Chinese |
|---|---|---|---|---|---|
| M01 - M05 | *Boletus edulis* | Penny Bun, cep, porcino | Karljohan, stensopp | белый гриб, боровик | 美味牛肝菌 |
| M06 - M09 | *Leccinum aurantiacum, Leccinum versipelle, Leccinum vulpinum, Leccinum albostipitatum* | red-capped scaber stalk | Aspsopp, Tegelsopp, Tallsopp | подосиновик, красноголовик | 橙黄疣柄牛肝菌 |
| M10 - M12b | *Amanita muscaria* | fly agaric, fly amanita | Röd flugsvamp | мухомор красный | 毒蝇伞 |
| M13 - M14 | *Russula paludosa, Russula sanguinaria, R. aurea, Russula emetica, Russula xerampelína, R. faginea* | brittlegill, bloody brittlegill, gilded brittlegill, sickener, rosey russula, emetic russula, vomiting russula | Storkremla, Blodkremla, Giftkremla, Guldkremla | красная сыроежка, сыроежка болотная, с. буреющая, сыроежка кроваво-красная, с. жгуче-едкая, с. рвотная | 沼泽红菇 |
| M15 - M16 | *Leccinum scabrum* | birch bolete , rough-stemmed bolete, scaber stalk | Björksopp | подберёзовик | 褐疣柄牛肝菌 |
| M17 | *Macrolepiota procera* | parasol mushroom | Stolt fjällskivling | гриб-зонтик высокий, гриб-зонтик пёстрый | 高大環柄菇 |
| M18 - M18b | *Stropharia caerulea, Stropharia cyanea, Stropharia aeruginosa, Psilocybe cyanea* | blue roundhead, blue-green stropharia | Blågrön kragskivling, Ärggrön kragskivling | строфария небесно-синяя, строфария сине-зелёная | 銅綠球蓋菇 |
| M19 - M19c | *Collybia nuda, Lepista nuda, Clitocybe nuda* | blewit, wood blewit | Blåmusseron | рядовка фиолетовая | 紫丁香蘑 |
| M20 - M20b | *Paxillus involutus, Paxillus rubicundulus* | brown roll-rim, common roll-rim | Pluggskivling, Alpluggskivling | свинушка тонкая, свинушка ольховая | 卷缘桩菇 |
| M20c - M20e | *Lactarius turpis, Lactarius necator; Paxillus involutus* | ugly milk-cap and brown roll-rim, common roll-rim | Pluggskivling och Svartriska | свинушка тонкая и чёрный груздь,чернушка | 醜乳菇 和 卷缘桩菇 |
| M21 - M21b | *Hydnum repandum* | hedgehog mushroom, sweet tooth | Blek taggsvamp | ежовик жёлтый | 美味齿菌 |
| M22 - M23 | *Craterellus tubaeformis* | yellowfoot, funnel chanterelle, winter chanterelle | Trattkantarell | лисичка трубчатая, лисичка ворончатая | 管形喇叭菌 |
| M24 - M24b | *Armillaria mellea, Armillaria bulbosa, A. gallica* | honey fungus | Honungsskivling, Klubbhonungsskivling | опёнок осенний, опёнок настоящий, опёнок толстоногий | 蜜環菌 |
| M25 - M25b | *Pholiota squarrosa* | shaggy scalycap, shaggy Pholiota | Fjällig tofsskivling | чешуйчатка обыкновенная, чешуйчатка ворсистая | 翘鳞伞 |
| M26 - M26b | *Albatrellus ovinus* | sheep polypore | Fårticka | трутовик овечий | 绵地花孔菌 |
| M27 | *Sparassis crispa* | cauliflower fungus | Blomkålssvamp | спарассис курчавый, спарассис кудрявый | 繡球菌 |
| M28 - M30 | *Ramaria sp., Ramaria flava, Ramaria formosa, Ramaria aurea, Ramaria pallida; Clavulinopsis sp.; Calocera viscosa* | yellow coral mushroom, pinkish coral mushroom, yellow-tipped coral fungus, pink coral fungus; coral mushroom; Clavulinopsis; yellow stagshorn | Gul fingersvamp, Lömsk fingersvamp, Blek fingersvamp; Clavulinopsis; klibbfingersvamp, hjorthornssvamp | рамария жёлтая, медвежья лапка, рогатик жёлтый, грибная лапша, оленьи рога, рамария красивая; клавулинопсис; калоцера клейкая | 珊瑚菌, 刷把菌,<br>疣孢黄枝瑚菌 |
| M31 - M31b | *Sarcodon imbricatus, Sarcodon squamosus* | shingled hedgehog, scaly hedgehog | Fjällig taggsvamp, Motaggsvamp | ежовик пёстрый, ежовик чешуйчатый | 翘鳞肉齿菌 |
| M32 - M32b | *Phellodon connatus, P. melaleucus, P.niger* | grey tooth; black tooth | Svartvit or Svart taggsvamp | ежовик сросшийся | 黑白栓齿菌, 黑栓齿菌 |
| M33 - M34 | *Fomes fomentarius* | tinder fungus, tinder conk, tinder polypore, hoof fungus, ice man fungus | Fnösticka | трутовик настоящий | 木蹄層孔菌 |
| M35 | *Phellinus igniarius* | willow bracket, false tinder conk | Eldticka | трутовик ложный | 发火木层孔菌 |
| M36 - M37 | *Fomitopsis pinicola* | red-belted conk | Klibbticka | трутовик окаймлённый | 松生擬層孔菌 |
| M38 | *Phaeolus schweinitzii* | velvet-top fungus, dyer's polypore, dyer's mazegill, pine dye polypore | Grovticka | трутовик Швейница | 栗褐暗孔菌 |
| M39 | *Trichia decipiens* | salmon-eggs | Gul ullklubba | трихия обманчивая | 長尖團毛黏菌 |
| M40 | *Fuligo septica* | scrambled egg slime, flowers of tan, jasmine mold | Trollsmör | земляное масло, муравьиное масло, фулиго гнилостный | 煤绒菌 |
| M41-M41c | *Termitomyces yunnanensis* | termite mushroom, chinese termitomyces | ? | юньнаньский термитомицес | 鸡枞菌 |
| M42-M42a | *Lactifluus volemus, Lactarius volemus* | weeping milk cap, bradley | Mandelriska | груздь красно-коричневый | 多汁乳菇, 红奶浆菌 |
| M43-M43a | *Russula viresens* | green-cracking russula, quilted green russula | Rutkremla | сыроежка зеленоватая | 青头菌, 变绿红菇 |
| M44 | *Neoboletus luridiformis, Boletus luriformis discolor* | scarletina bolete, red foot bolete, dotted stem bolete | Blodsopp | дубовик крапчатый | 紅柄牛肝菌 |
| M45-M45a | *Turbinellus matijun, Gomphus matijun* | matijun | ? | гомфус матиджан | 馬蹄均 |
| M46 | *Cantharellus applanatus* | flat chanterelle | ? | лисичка плоская | 鸡油菌 |
| M47-M47a | *Rugiboletus extremiorientalis* | rugiboletus | ? | обабок дальневосточный | 皱牛肝菌属, 皱牛肝菌 |
| M48 | *Suillus granulatus* | weeping bolete, granulated bolete | Grynsopp | маслёнок летний, маслёнок зернистый | 栗壳牛肝菌 |
